# A data assimilation method to track time-varying changes in the excitation-inhibition balance using scalp EEG

**DOI:** 10.1101/2022.12.16.520838

**Authors:** Hiroshi Yokoyama, Keiichi Kitajo

## Abstract

Recent neuroscience studies have suggested that controlling the excitation and inhibition (E/I) balance is essential for maintaining normal brain function. However, while control of time-varying E/I balance is considered essential for perceptual and motor learning, an efficient method for estimating E/I balance changes has yet to be established. To tackle this issue, we propose a new method to estimate E/I balance changes by applying neural-mass model-based tracking of the brain state using the Ensemble Kalman Filter. In this method, the parameters of synaptic E/I gains in the model are estimated from observed electroencephalography (EEG) signals. Moreover, the index of E/I balance was defined by calculating the ratio between synaptic E/I gains based on estimated parameters. The method was validated by showing that it could estimate E/I balance changes from human EEG data at the sub-second scale, indicating that it has the potential to quantify how time-varying changes in E/I balance influence changes in perceptual and motor learning. Furthermore, this method could be used to develop an E/I balance-based neurofeedback training method for clinical use.

## Introduction

A balance between synaptic excitation and inhibition (E/I) plays a crucial role in the neural mechanisms underlying social behavior, neuropsychiatric disorder-related symptoms, perceptual performance, sleep function, and synaptic homeostasis. For example, an animal study has shown that synaptic E/I imbalance within the prefrontal cortex leads to autistic-like social dysfunction [1]. In the context of human neuroscience, studies using concurrent transcranial magnetic stimulation and electroencephalography (TMS-EEG) recordings have reported evidence of a relationship between cortical GABAergic inhibition/glutamatergic excitation and psychiatric disorders [2, 3]. Therefore, the disruption of E/I balance in the human brain could also lead to social dysfunction. With the recent advances in neurotransmitter imaging using magnetic resonance spectroscopy (MRS), the in-vivo E/I balance can be evaluated within the intact human brain. For instance, two human studies suggested the essential role of controlling E/I balances for the consolidation of perceptual and motor memory [4, 5]. Both studies indicated that the excitation-dominant changes in E/I balance resulting from down-regulation of GABA concentration played a role in facilitating plastic changes during perceptual and motor learning. In addition, one of these studies [4] revealed that the continued training of a skill after performance improvement (i.e., overlearning) led to the enhancement of inhibition-dominant changes in E/I balance and consolidation of synaptic plasticity, resulting in the stabilization of perceptual learning. Therefore, it is important to track time-varying changes in E/I balance to understand the functional mechanisms of perceptual learning. However, the temporal dynamics of changes in E/I balance during perceptual learning are still unknown since temporal changes in in-vivo neurotransmitters are difficult to track on a moment-to-moment basis because it takes about 10 min for MRS to acquire data from a region of interest. To address these issues, it is important to establish a method that can measure temporal changes in E/I balance in the intact human brain on a moment-to-moment basis.

The changes in E/I balance can be considered as changes in synaptic current mediated by glutamatergic excitation and GABAergic inhibition. Thus, E/I balance changes would lead to changes in the dynamics of postsynaptic potentials, local field potentials, and scalp EEG. We predicted that the activity in human scalp EEG would reflect the temporal dynamics of E/I balance. In fact, recent TMS-EEG studies suggested that TMS-evoked potentials (TEP) in scalp EEG reflected both GABAergic and glutamatergic mediated functions [2, 3, 6, 7]. However, to our knowledge, an efficient method to directly quantify the E/I ratio using only observed EEG signals has not been established. To this end, we propose a new model-based method to estimate temporal changes in E/I balance from observed EEG signals by applying neural-mass (NM) model-based tracking of the brain state using the Ensemble Kalman Filter (EnKF). The EnKF [8, 9] was developed to assimilate the nonlinear dynamical models with observed data. In the field of control theory, the nonlinear Kalman filter scheme (including the EnKF) is applied not only to predict time-series data but also to estimate model parameters. Furthermore, computational studies indicated that the time-evolving dynamics of EEG signals (EEGs) can be formulated as a nonlinear dynamical system, such as in the NM model [10], which models interactions between pyramidal cells, excitatory interneurons, and inhibitory interneurons. Based on the above, we assumed that, by directly assimilating the NM model to observed human scalp EEGs, the temporal changes in E/I balance could be quantified according to the NM model parameters representing the E and I synaptic gains estimated from the observations. Our proposed method to evaluate the in-vivo E/I balance in the brain has the potential to be an effective way of overcoming the above-mentioned limitation of the prior method.

Here, we introduce an overview of our proposed method. In this method, to predict the time-series of the observed EEGs and estimate the NM model parameters in parallel, the single-channel time-series of the EEGs are sequentially fitted to an NM model [10] using variational Bayesian noise adaptive constrained EnKF (vbcEnKF) on a sample-by-sample basis. Using the estimated parameters reflecting the E and I synaptic gains, the temporal changes in the ratio of E and I postsynaptic activities are evaluated as a putative index of the model-based E/I balance (mE/I ratio).

To validate our proposed method, two datasets were analyzed. First, we applied our proposed method to the synthetic data generated by the NM model with known temporal changes in the model parameters. By doing so, we confirmed that the estimated model parameters obtained using our method were consistent with the exact model parameters. Next, we applied our method to an open dataset of experimental EEG signals measured from healthy humans while sleeping [11]. In this way, we tested whether our proposed method could detect the time-varying changes in E/I balance during sleep from observed EEG signals.

## Results

### Proposed method

In our proposed method, we evaluate the time-varying changes in the E/I ratio according to the parameters reflecting the E and I synaptic gains in the NM model (see the description for the NM model parameters *A* and *B* in Methods section). The single-channel time series of observed EEGs are sequentially fitted to the model based on the data assimilation scheme using vbcEnKF. We provide an outline of our proposed method, especially concerning how the time-varying changes in the E/I balance are estimated from observed EEGs using vbcEnKF. For more mathematical details on vbcEnKF, please see the Methods section and Supplementary Information.

In this study, we assumed that the observed EEGs *y*_*t*_ could be formulated as the following state-space form of the NM model:

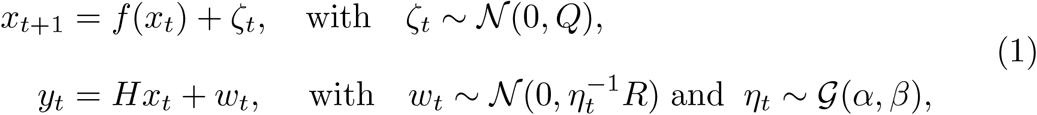

where *x*_*t*_ = {**v**_*t*_, *θ*_*t*_}^*T*^ stands for the state variables containing the NM model variables *v*_*t*_ and the model parameters *θ*_*t*_. The variable *H* indicates the linear coefficient for the observation model. The function *f* (*x*_*t*_) is formulated as 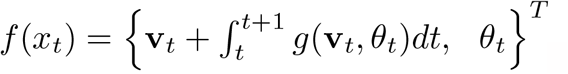, where the term 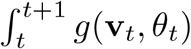 indicates the numerical integral of the NM model *g*(**v**_*t*_, *θ*_*t*_), which is implemented by the fourth-order Runge-Kutta method with time-step Δ*t*. In our method, Δ*t* is set to the sampling interval of the observed data. Detailed descriptions of the NM model are provided in the Methods section. The variable *ζ*_*t*_ represents the state noise that follows the Gaussian distribution N(0, *Q*). The variables *w*_*t*_ and *η*_*t*_ are the observation noise and noise scaling factor that follow the Gaussian distribution 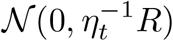 and gamma distribution G(*α, β*), respectively. *Q* and *R* are the fixed covariance parameters for the state and observation noise, respectively. In our method, the state *x*_*t*_ and observation *y*_*t*_ follow Gaussian distributions as *x*_*t*_ ∼ 𝒩 (*x*_*t*_, *P*_*t*_) and 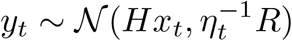, respectively. To estimate the state *x*_*t*_ and the noise scaling factor *η*_*t*_ from the observation *y*_*t*_, the following equation based on Bayesian theory can be applied:

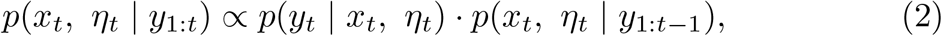

where *y*_1:*t*_ indicates the observation set as {*y*_1_, *y*_2_, …, *y*_*t*_}. In such a case, if *x*_*t*_ and *η*_*t*_ are independent, the joint posterior distribution *p*(*x*_*t*_, *η*_*t*_ | *y*_1:*t*_) in equation (2) can be solved by minimizing the Kullback–Leibler (KL) divergence between the variational Bayesian approximation *q*(*x*_*t*_)*q*(*η*_*t*_) and the exact distribution *p*(*x*_*t*_, *η*_*t*_ | *y*_1:*t*_) [12]:

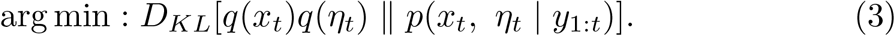

In our proposed method, by combining the solution to equation (3) [12–16] and the constrained EnKF-based [17–20] state estimation scheme (which we named vbcEnKF), we can derive an estimation rule of the state *x*_*t*_ = {**v**_*t*_, *θ*_*t*_}^*T*^ and noise scaling factor *η*_*t*_ (see the Methods section and Supplementary Information for more details on vbcEnKF). On the basis of these solutions, the time-varying changes in the model variable **v**_*t*_ and parameter *θ*_*t*_ can be estimated from the observed EEGs *y*_*t*_ on a sample-by-sample basis. Since the estimated *θ*_*t*_ is containing the parameters representing the E and I synaptic gains, i.e., *A*(*t*) and *B*(*t*) in the NM model (see the Methods section for more details), the time-varying changes in the E/I balance can be quantified by calculating the ratio of these parameters. Therefore, the evaluation index of the E/I ratio (mE/I ratio: mE*/*I(*t*) = *A*(*t*)*/*(*A*(*t*) + *B*(*t*))) can be calculated from the estimated *θ*_*t*_. Using this index, we can directly estimate the time-varying changes in the E/I ratio from the observed EEG signals. A schematic of this proposed method is shown in Fig. 1.

**Fig. 1.**
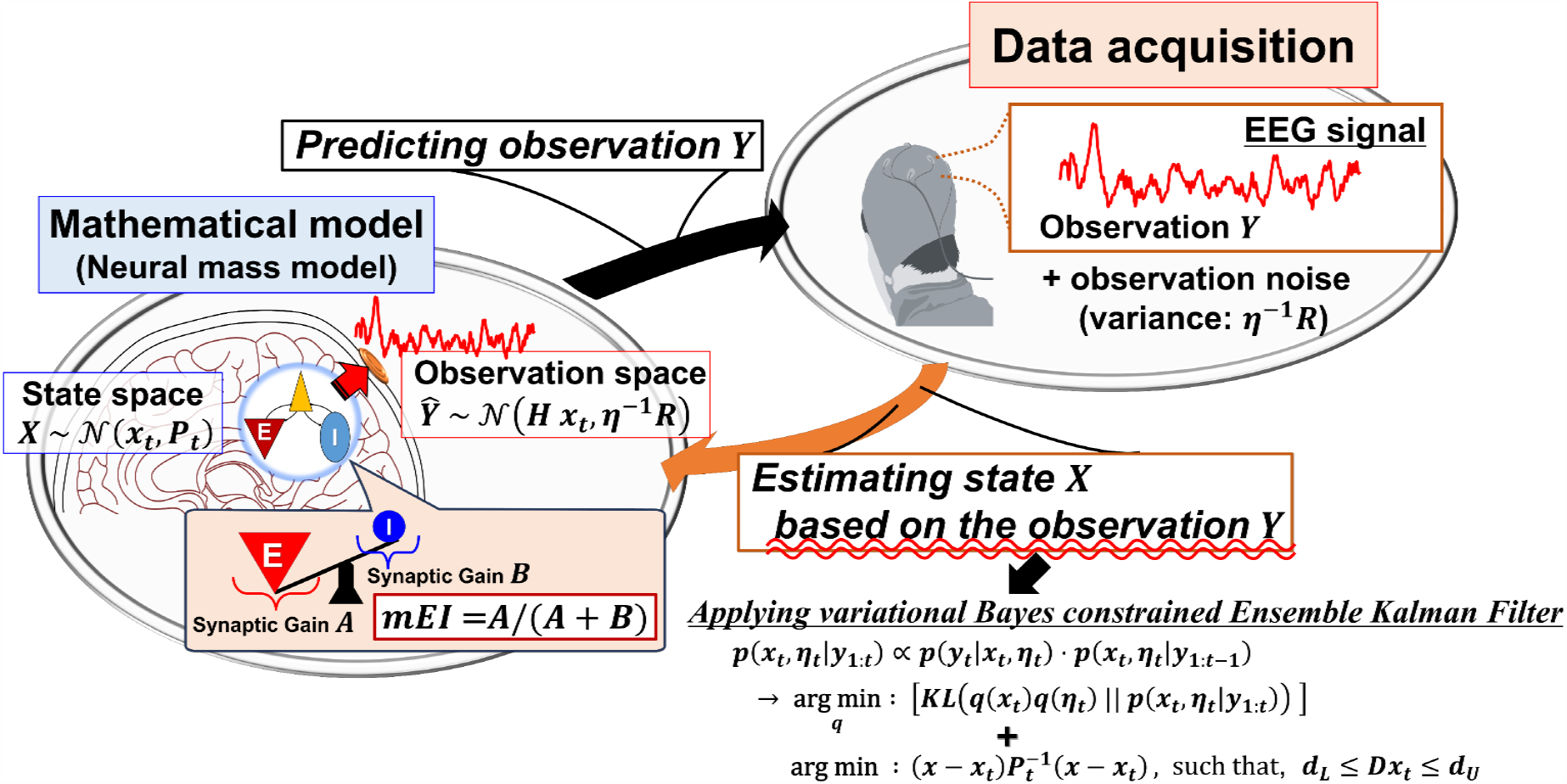
Overview of the proposed method. The single-channel time-series data from observed EEGs are sequentially predicted by a neural-mass (NM) model using the variational Bayesian noise adaptive constrained Ensemble Kalman Filter (vbcEnKF), and the state variable *x*_*t*_, which is containing the parameters *A* and *B* representing the E and I synaptic gains in the model, is estimated from the observed EEGs *y*_*t*_. Based on our assumption that the changes in the excitation-inhibition (E/I) balance are reflected by changes in the estimated model parameters *A* and *B*, we propose the model-based evaluation index for the E/I ratio: (mE/I ratio = *A/*(*A* + *B*)).

### Verification with numerical simulation

To confirm whether our proposed method can correctly estimate the temporal changes in model parameters from the observed data, we applied our method to synthetic data generated by an NM model with known parameter changes.

Moreover, as described in the Methods section, our proposed method applied the combination approach of EnKF with a variational Bayesian noise adaptive algorithm [12–16] to avoid reduction in estimation accuracy caused by the non-stationarity of the observation noise. Therefore, we also evaluated whether the estimated covariance of observation noise was consistent with exact noise covariance containing the synthetic EEG signal. In addition, since the EnKF is based on the sequential Monte Carlo (sMC) method for Bayesian probability estimation, both the initial seed value and ensemble size for the random sampling should affect the error of state estimation. Therefore, to compare the prediction error according to the ensemble size and initial random seed of sMC sampling, we selected ensemble sizes from 40 to 500 with 20 steps and applied these to the same synthetic data 50 times with a different initial random seed for each ensemble size (see the Methods section for a more detailed description of simulation settings).

The results of this validation are shown in Figs. 2 and 3. Fig. 2 shows examples of state estimation results with *N*_*ens*_ = 40 and 200. While the predicted EEG time series was similar to the exact synthetic EEG between both ensemble size conditions (Fig. 2a, d), the estimation of the mE/I ratio tended to be accurate with respect to the increase of ensemble size (Fig. 2b, e). Moreover, the estimation of the model parameters also tended to be accurate depending on the increase of ensemble size (Fig. 2c, f). To quantitatively reveal the prediction skill according to the ensemble size in our method, we then assessed the accuracy of EEG signal prediction and observed the estimated noise covariance (Fig. 3). As shown in Fig. 3a, the prediction error of the EEG signal, based on the mean absolute error score, showed a tendency toward decreasing the error with an increase in the ensemble size *N*_*ens*_. This tendency was also found in the estimation results of observation noise covariance (Fig. 3b). Moreover, estimated noise covariance converged to exact noise covariance with *N*_*ens*_ ≥ 200.

**Fig. 2.**
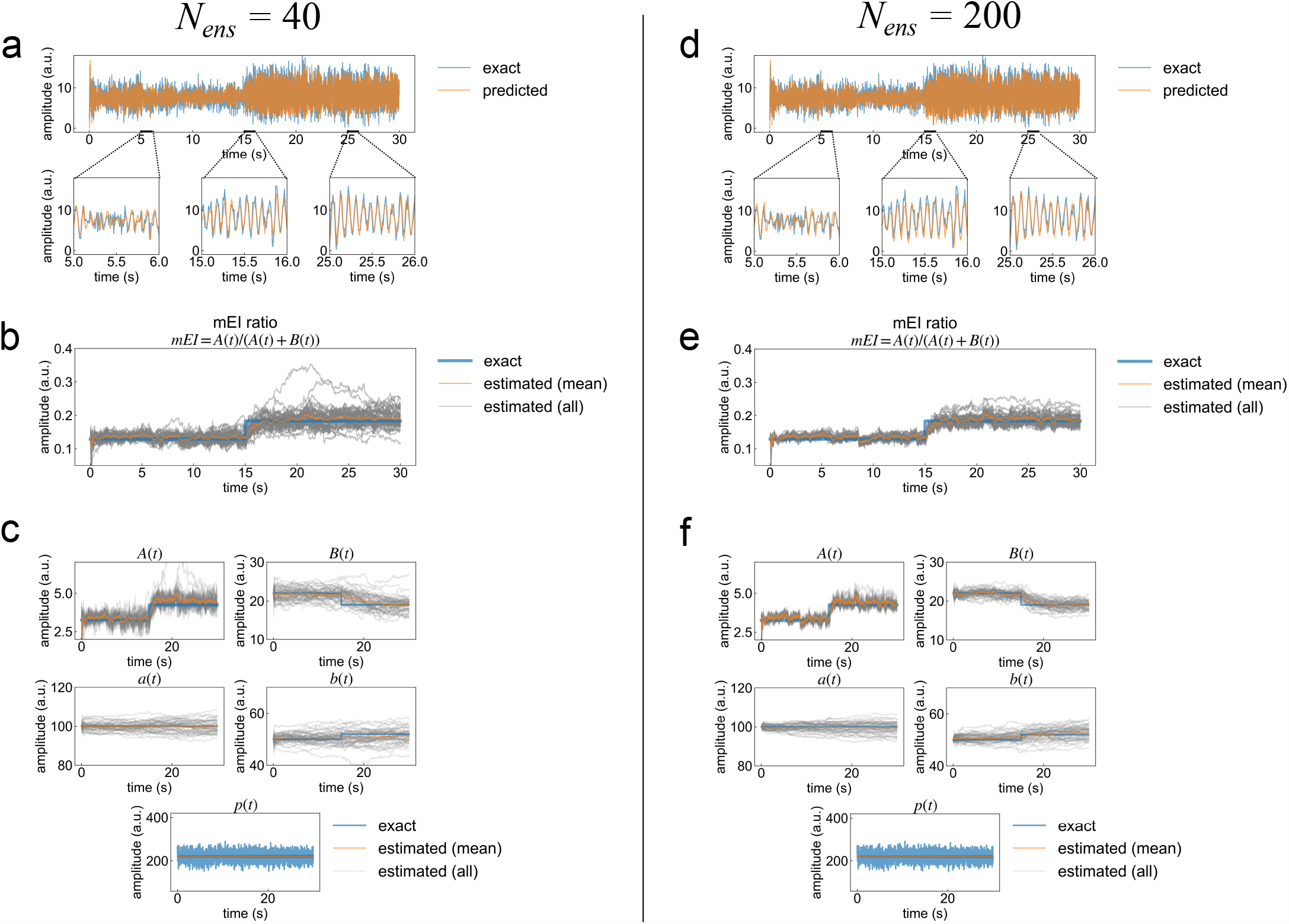
Prediction of EEG time-series data and estimation of unobservable model parameters using synthetic EEG data. a, d) Comparison between the original EEG data and typical predicted EEG results obtained from the first trial out of 50 trials with ensemble sizes of *N*_*ens*_ = 40 and 200, respectively. The upper panels in (a) and (d) show the whole time series of synthetic and predicted EEG. The lower three panels in (a) and (d) are enlarged views of the prediction results for each time interval. Blue and orange lines indicate the exact and predicted values of the EEGs. b, e) Estimations of the mE/I ratio obtained from 50 trials with ensemble sizes of *N*_*ens*_ = 40 and 200, respectively. c, f) Estimations of the five parameters, A,a,B,b, and p, obtained from 50 trials with ensemble sizes of *N*_*ens*_ = 40 and 200, respectively. Blue and orange lines in (b)––(f) indicate the time series of exact values and the mean of the estimations. Gray color lines in (b)––(f) indicate the estimated value of each parameter obtained from 50 trials.

**Fig. 3.**
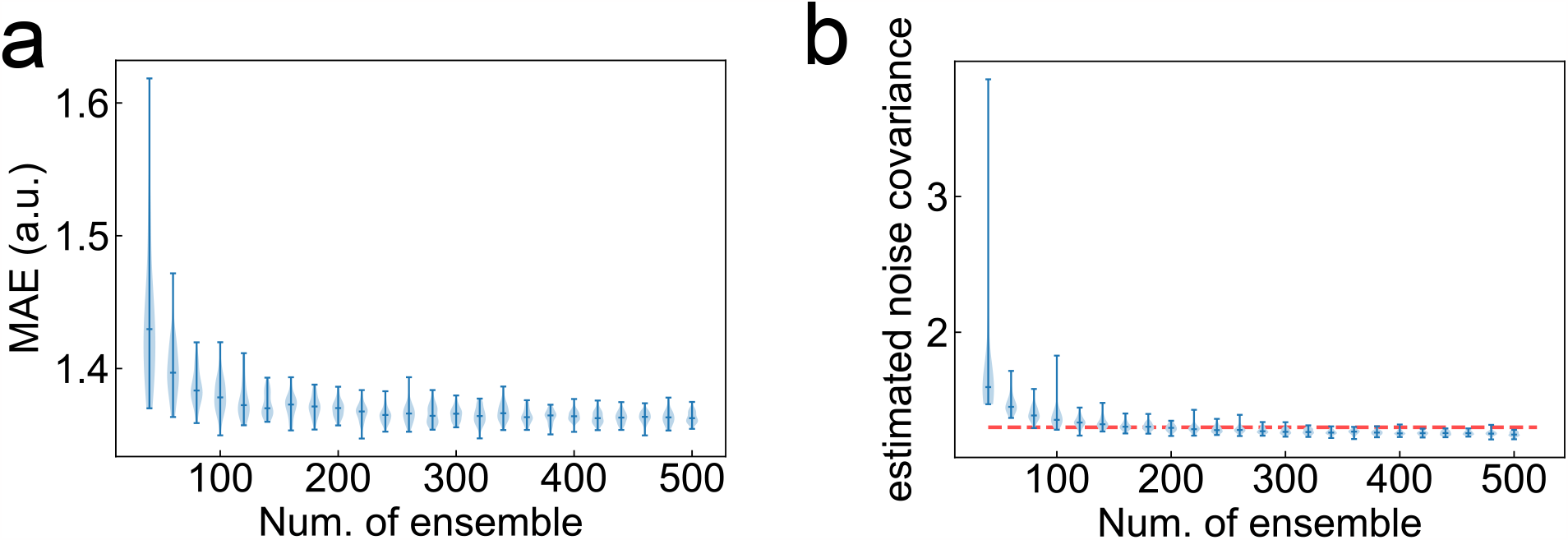
Effects of ensemble size on estimation accuracy. a) Violin plots of prediction error scores as a function of the number of ensembles. The probability densities in these violin plots were estimated using a kernel density estimation method based on the samples obtained from 50 trials. The error bar indicates the maximum and minimum value of samples. The middle line of the error bar shows the median value of the samples. b) Corresponding estimated observation noise covariance as a function of the number of ensembles. The red-dot line indicates the exact noise covariance of synthetic EEG data.

In summary, the proposed approach was able to predict the model states and estimate parameters, and *N*_*ens*_ ≥ 200 was required to guarantee an accurate prediction with a smaller prediction error.

### Sleep EEG analysis

To confirm the neurophysiological validity of our proposed method, we applied it to real human EEG data obtained from Kemp et al. (2000) [11]. The main aim of this validation was to assess whether our proposed method can track the sleep-dependent changes in the E/I balance between NREM/REM stages, such as those that are reported in both animal and human studies [21, 22]. Moreover, in the following analysis, we also compare the evaluation results between a prior method of EEG-based E/I ratio analysis [23] and our proposed method. The dataset from Kemp et al. (2000) [11] includes EEG recorded during day-night periods (over around 20 hours) from healthy participants at their home. The sleep-stage markers were scored manually by referring to recorded signals, and a more detailed description of the task settings is provided in the original paper [11]. In general, when sleeping, EEG oscillations between the delta and lower beta frequency band change depending on the sleep stages, such as nonrapid eye movement (NREM) or rapid eye movement (REM) sleep. Moreover, recent animal and human studies in sleep neurophysiology have suggested that sleep regulates E/I balance [21, 22]. In particular, Tamaki et al. (2020) [22] reported that the changes in the E/I balance in the early visual area are sleep-stage dependent. Using simultaneous magnetic resonance spectroscopy (MRS), the authors found that modulation of the E/I balance during sleeping played a role in stabilizing visual learning. Therefore, we applied our method to human sleep EEG data to determine whether it could track changes in the E/I balance according to the NREM/REM sleep stage, and compared the results from our method with those obtained from a prior EEG-based E/I balance estimation method [23] (hereafter called the E/I slope method. See the Methods section for details). For this analysis, we used EEG data from 19 healthy participants who showed typical transitions between the sleep stages during the first NREM (Stage 1, Stage 2, and Stage 3/4) and REM periods. Moreover, in the following validation tests, we used an EEG interval from the first 20-minute period before sleep onset to the end of the first REM period for each participant. In the following analysis, the interval from the first 20-minute period before the sleep onset was defined as the awake period.

A typical result of the time-series prediction of EEG in one participant (ID: 4032) is shown in Fig. 4. Our method correctly tracked the time-evolving dynamics of the original EEG signal within the sleep and awake intervals (Fig. 4 b–d). The results of the time-varying changes in the mE/I ratio estimated by our proposed method for two typical participants are shown in 5 a, b. In our method, while the estimated E/I ratio in the NREM period increased (i.e., synaptic gain of excitatory interneurons tended to be dominant during NREM periods), the value in the REM period decreased (i.e., synaptic gain of excitatory interneuron tended to be suppressed during REM periods). This result is consistent with the experimental results of an MRS study conducted by Tamaki et al. (2020) [22]. Moreover, the mE/I ratio estimated by our proposed method in the awake period was also smaller than that in the NREM period, and this was seen in both participants’ data (see Fig. 5a, b). A similar tendency was reported in an animal study [21]. In contrast to the estimations of the E/I ratio by our method, the E/I slope method proposed by Gao et al. (2017) [23] could not identify sleep-dependent changes in either dataset (Fig. 5 c, d).

**Fig. 4.**
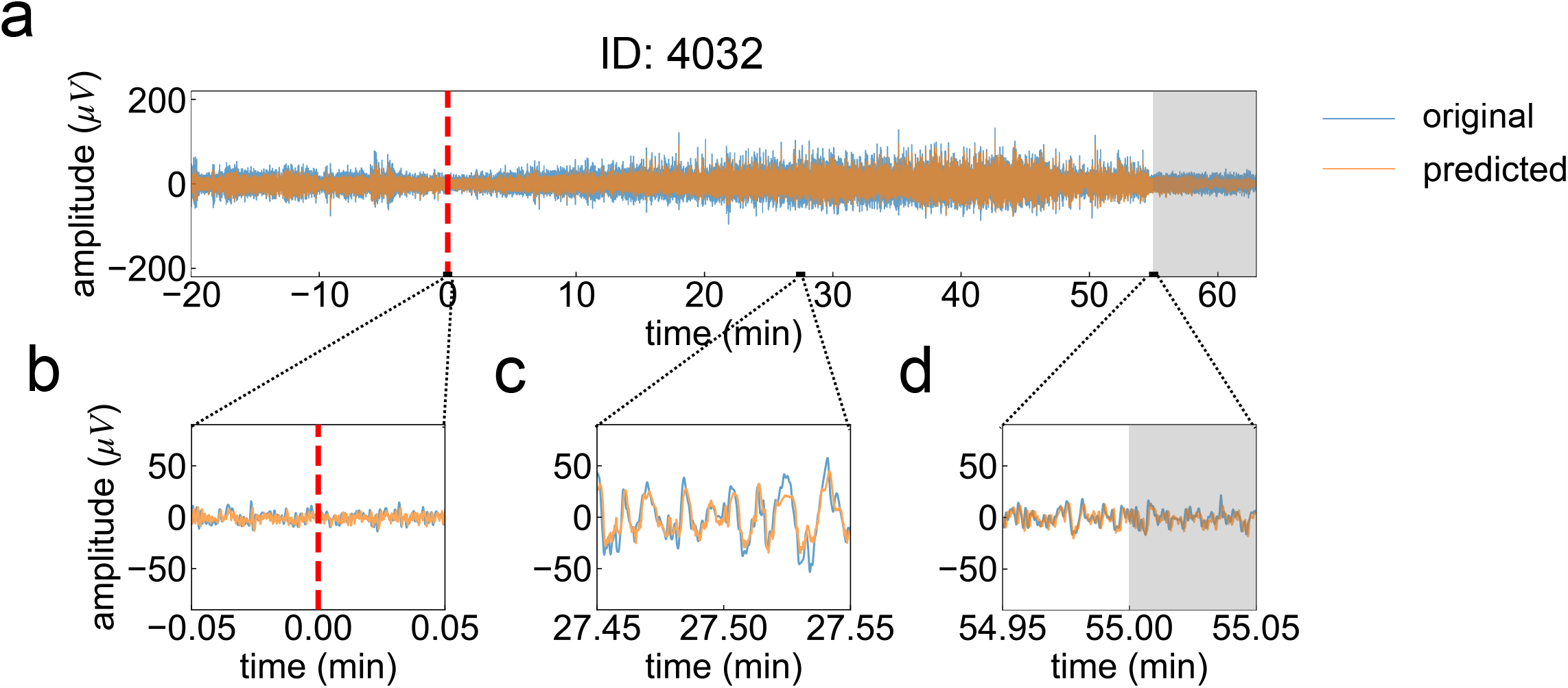
Typical examples of predicted EEG signals. a) Prediction result of an EEG signal. Blue and orange lines indicate the time series of observed and predicted EEG signals, respectively. On the x-axes, the position at time = 0 (min) indicates the sleep onset (onset of NREM period). The gray shaded area indicates the REM sleep period. b–d) Enlarged views of predicted signals with a 0.1 min time window for three-time intervals around sleep onset (b), middle of NREM (c), and around the end of NREM (i.e., the onset of REM) periods (d). The sleep stages in this dataset were manually determined by well-trained technicians [11].

**Fig. 5.**
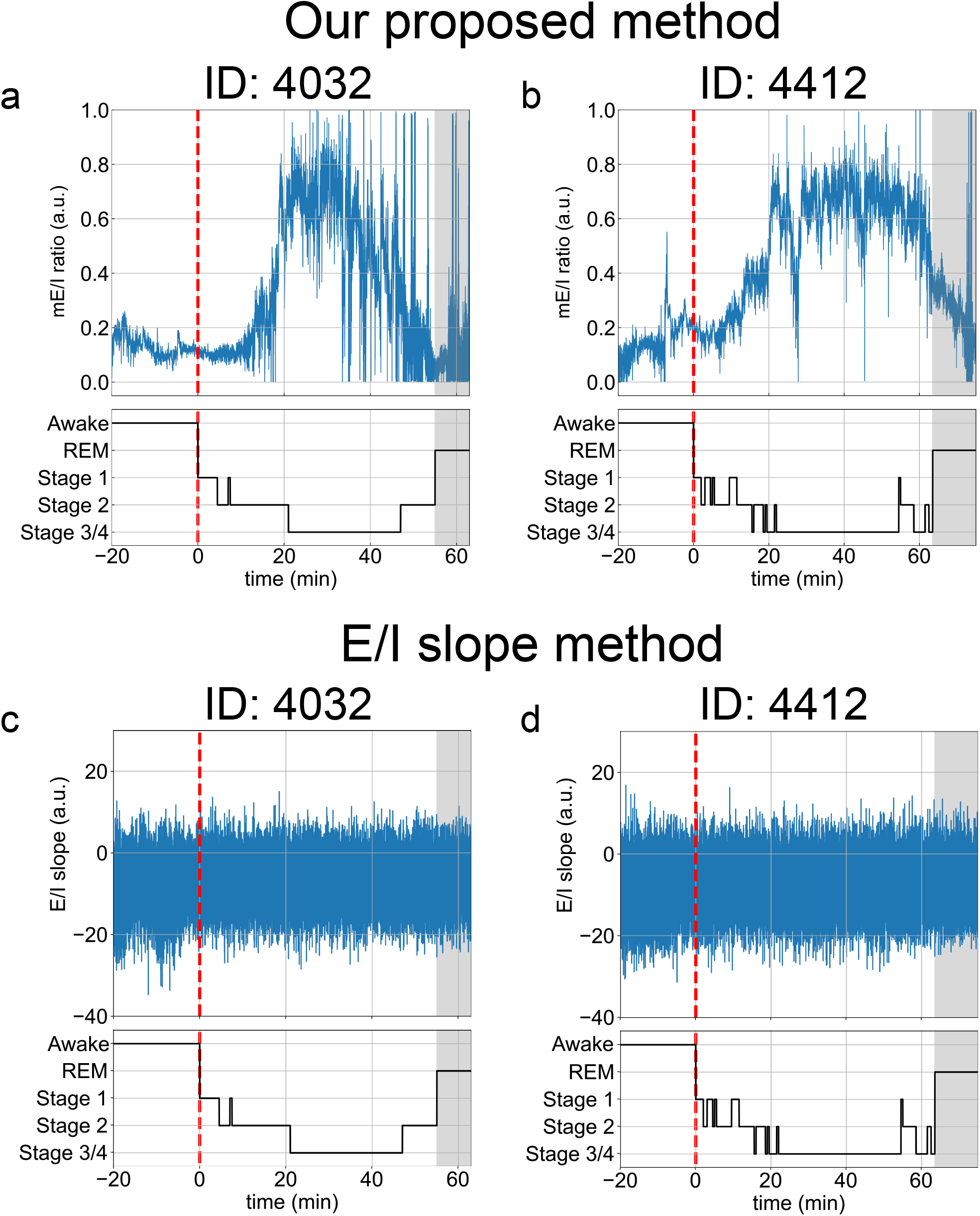
Comparison of the time-varying changes in the E/I ratio between our proposed method and the E/I slope method. a, b) Typical examples of the results of our proposed method for two participants. The upper panels of (a) and (b) show the time series of the mE/I ratio estimated from the observed EEG signals by our proposed method. Blue lines indicate the time series of the mE/I ratio. The lower panels in (a) and (b) show the time profiles of sleep-stage changes. Red dotted lines indicate the sleep onset (onset of NREM period). Gray shaded areas indicate the REM period. c, d) Typical examples of the results of the E/I slope method for the same two participants.

Next, to address the statistical validity of the E/I results, the time-averaged E/I ratio of each method (our proposed method and the E/I slope method) was evaluated for each sleep stage (Awake, State 1, State 2, State 3/4, and NREM) for all 19 participants (Fig. 6). Then, for each E/I estimation method, the group data of these time-averaged E/I ratios were subjected to a Kruskal-Wallis test, which is a rank-based nonparametric version of a one-way ANOVA. This Kruskal-Wallis test showed that the time-averaged E/I ratios obtained by our method (i.e., time-averaged mE/I score) were significantly different across distinct sleep stages (h-statistics *H* = 32.1325, *p <* 0.0001, effect size *η*^2^ = 0.3126), whereas those obtained by the E/I slope method showed no significant differences across distinct sleep stages (h-statistics *H* = 9.1670, *p <* 0.0570, effect size *η*^2^ = 0.0574). Furthermore, in the result of our proposed method, by subjecting the evaluated value to the Dunn’s multiple comparison test (two-tailed test), which is a rank-based method for post-hoc comparisons, the time-averaged mE/I ratio in Stage 3/4 (i.e., NREM Stage 3/4) was significantly higher than those in the Awake, Stage 2, and REM periods (Stage 3/4 vs. Awake,

**Fig. 6.**
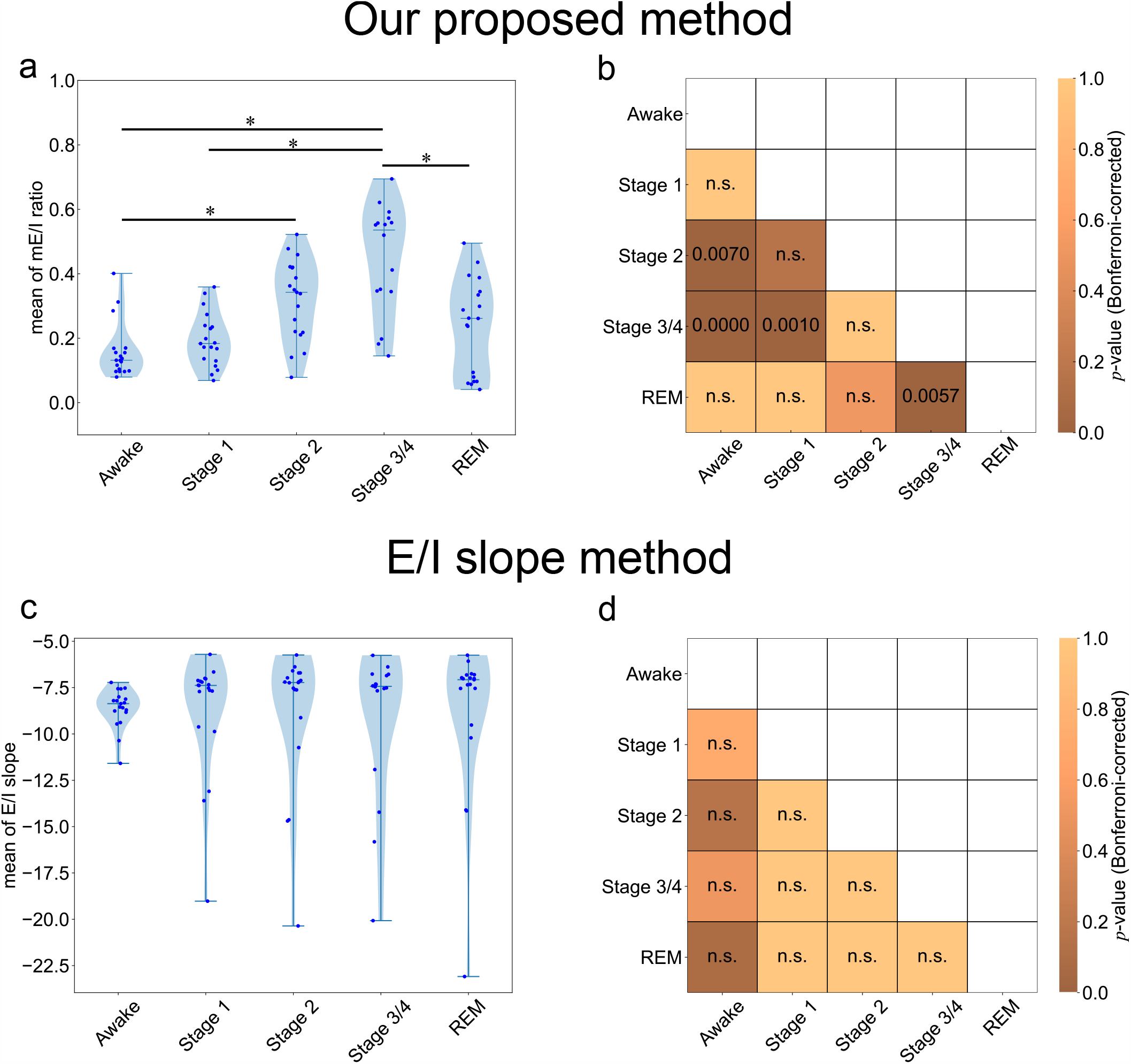
Comparison of the sleep-state dependent changes in the estimated E/I ratio between our proposed method and the E/I slope method. a, c) The violin plots were obtained from the samples of the time-averaged E/I ratios of 19 participants for each sleep stage, evaluated by our proposed method and the E/I slope method, respectively. The error bars indicate the maximum and minimum values of samples. The middle lines of the error bars show the median values of the samples. The blue dots indicate the samples of the time-averaged mE/I ratios for each participant. Asterisks (*) indicate *p <* 0.01 according to Dunn’s multiple comparison test with Bonferroni correction. b, d) Statistical results of the multiple comparisons of each pair of E/I ratios by sleep stage for our proposed method and the E/I slope method, respectively. *p*-values were adjusted using the Bonferroni method.

*p <* 0.0001; Stage 3/4 vs. Stage 2, *p* = 0.001; Stage 3/4 vs. REM, *p* = 0.0057; *p*-values were corrected using the Bonferroni method; Fig. 6a, b). Moreover, the mE/I ratio in Stage 2 (i.e., NREM Stage 2) was significantly higher than in the awake period (*p* = 0.007). Other paired comparisons of sleep stages showed no significant differences for the time-averaged mE/I ratios (Awake vs. Stage 1, *p* = 1.00; Awake vs REM, *p* = 1.00; Stage 1 vs. Stage 2, *p* = 0.16; Stage 1 vs. REM, *p* = 1.00; Stage 2 vs. Stage 3/4, *p* = 1.00; Stage 2 vs. REM, *p* = 0.52). In contrast to the results of our method, as shown in Fig. 6 c and d, no significant difference was found in the results of the time-averaged E/I slope values (prior method) for any pair of sleep stages.

In summary, these results indicated that changes of the mE/I ratio estimated by our proposed method using observed EEG signals were consistent with the experimental evidence reported by prior studies [21, 22].

## Discussion

In this study, we proposed a new approach to track the time-varying changes in E/I balance from observed EEG data on a sample-by-sample basis. Using both numerical and empirical neurophysiological data, our validation results supported our assumptions, as follows: (1) Temporal changes in the E/I ratio caused by synaptic current changes are reflected in the dynamics of scalp EEG, and (2) temporal changes in the E/I ratio could be evaluated using estimated parameters in the NM model with the EnKF scheme. Moreover, sleep stage-dependent changes in the mE/I ratio estimated from sleep EEG data (see Figs. 5 and 6) were consistent with previous experimental MRS results reported by Tamaki et al. (2020) [22]. Based on these previous results and our own, we suggest that the proposed method could be applied to human EEG to reveal the sleep-related mechanisms underlying changes in the E/I ratio. Moreover, while MRS cannot track time-varying changes in the E/I ratio because of its inherent measurement limitations, our method can estimate such time-varying changes using the proposed metric, i.e., the mE/I ratio. This difference in the temporal resolution for tracking E/I balance changes is the most important advantage of our proposed method.

In human neuroscience, two methods are used to measure E/I balance changes in the intact brain. The first one is TEP, which is based on TMS-EEG [24, 25]. Some experimental evidence obtained using this method has suggested that TEP changes recorded in human scalp EEGs reflect both GABAergic-mediated inhibitory and glutamatergic-mediated excitatory functions [6, 7]. Therefore, by evaluating TEP using TME-EEG methods, we can determine how intracortical E/I reflects the neural mechanisms underlying a neuropsychiatric disorder associated with E/I balance changes [2, 3]. TEP is assessed by calculating the TMS-triggered averaged EEG signals with TMS stimulation with around 2 s inter-pulse-intervals (IPI) over multiple trials [24, 26], and TMS-EEG-based evaluations of intracortical E/I require at least 60 trials to define the TEP component of EEG signals [26]. Therefore, TEP based cortical E/I evaluations are limited to measuring EEG signals at intervals of at least 2 min (2 s IPI × 60 trials ≒ 120 s) for the detection of E/I related EEG components. The second method employs in-vivo imaging of brain neural metabolites using MRS. As mentioned in the Introduction, a recent MRS study [4] suggested that overlearning leads to changes in the inhibition-dominant E/I balance to enhance stabilization of perceptual learning. In addition, another study [22] suggested that sleep-induced E/I balance changes play a role in consolidation of the memory of skill learning before sleep. However, since the MRS method cannot track time-varying changes in E/I balance due to the imaging scan time constraint (around 10 min to scan a region of interest [4, 22]), it is not clear how temporal changes in synaptic plasticity are involved in consolidating the novel perceptual skill. Taken together, TMS-EEG and MRS-based methodologies are cutting-edge measures for estimating the E/I balance from the in-vivo brain; however, both methods cannot track time-varying changes in the E/I balance from moment-to-moment. In contrast to these methods, the method proposed in this study can overcome these issues by directly estimating time-varying changes in the E/I ratio from moment-to-moment in the EEG data on a single-trial basis and on a sub-second scale. The originality of our proposed method lies in that the method can reconstruct unobserved features of the E/I ratio from observed EEG on a single-trial basis by applying data assimilation techniques. These advantages suggest that it could be used to advance understanding of how the synaptic plasticity changes, reflected in the time-course of E/I balance at the sub-second scale, are involved in perceptual skill and its enhancement during a learning task. The proposed method could find applications in neurorehabilitation. For instance, it could be used to develop a skill training method based on E/I balance-based neurofeedback for the neurorehabilitation of patients with severe motor disorders such as stroke.

Concerning previous studies on local field potentials (LFPs), one study proposed a time-frequency analysis-based approach to track temporal changes in E/I balance from LFP signals [23]. This study demonstrated that the slope of the log-log plot of the power spectral density functions between 30 and 50 Hz in LFPs (i.e., 1/*f* power law exponent in the gamma frequency band) was correlated with the E/I ratio. Therefore, the authors proposed a power spectral density-based E/I estimation method (the E/I slope method). Moreover, a recent study [27] applied E/I slope analysis to human EEG data and found a significant difference in the E/I balance between awake and REM states. Therefore, we applied this method (called the E/I slope method in this paper) as a prior method of EEG-based E/I balance estimation for comparisons with the results of our proposed method. As shown in 5c, d, and 6c, d, sleep-dependent changes in the E/I ratio were not observed when using the E/I slope method, unlike previous findings [27]. As mentioned in the Results section, no significant difference in the results of the E/I slope method were found, regardless of the sleep stages. The failure of the E/I slope method to detect sleep-stage changes in our dataset may be because gamma activity (*>* 30 Hz) in EEG tends to be compromised due to electromyogenic artifacts from cranial and ocular muscles. Moreover, since the sampling frequency of EEG signals in the dataset we used was 100 Hz (i.e., the meaningful frequency components of observed EEG in our dataset were below the 50 Hz), the signal of gamma band oscillations over 50 Hz would not be properly observed in this dataset. This could be considered why the E/I slope method cannot efficiently detect the sleep-dependent changes in E/I balance from EEG data. In practice, power spectra in the gamma band (especially those greater than 50 Hz) of our EEG data were weaker than those of lower frequency bands. In contrast, our proposed method can detect significant differences in the mE/I ratio between sleep stages, even for situations in which the E/I slope method cannot (Figs. 5a, b and 6a, b). Our method directly focused on frequency bands that indicate sleep-dependent oscillations under 20 Hz (i.e., delta, theta, and alpha oscillations) to evaluate the mE/I ratio from observed EEG data with the vbcEnKF scheme in the NM model. This contrast between our proposed and prior methods highlights the advantage of our method, in that it can estimate the E/I ratio from observed EEG data without relying on signal quality or sampling frequency. Moreover, by directly assimilating the observed EEG to the NM model, we can interpret how sleep-dependent EEG oscillations are caused by the synaptic balance between excitation and inhibition according to changes in the estimated E/I ratio. This is another advantage of our proposed method.

To the best of our knowledge, the first study to adapt data assimilation techniques in neuroscience research was conducted by Ullah et al. (2010) [28]. That study reported that an unobserved process of neuronal dynamics associated with cellular excitability during seizure can be reconstructed from a single observed membrane potential by assimilating the neuron model behavior to real observed data. In the past decade, some other studies have also used data assimilation techniques [28–31]. In particular, Kuhlmann et al. (2016) [30] proposed that the depth of anesthesia can be tracked by looking at the changes in predicted model parameters of the NM model from observed human EEG using the unscented Kalman Filter. These prior studies applied the state and parameter estimation approach using the nonlinear Kalman Filter or Bayesian filter for the data assimilation of neural data, as with our proposed method. However, those previous methods did not consider the effect of nonstationary observation noise on the accuracy of model parameter estimation in the Kalman Filter scheme. In addition, most neurophysiological models contain the parameter constraints to allow for stable dynamics of its model behavior; however, the prior methods did not consider this issue when assimilating the model prediction to observed data. Comparing our proposed method with those of the above-mentioned studies, through use of the vbcEnKF scheme, our method can track the time-evolving dynamics of observed signals while considering both the effect of nonstationary observation noise and parameter constraints in the model. As shown in the results of numerical simulations (see Fig. 3), since the vbcEnKF can correctly determine the observation noise scale with the ensemble size *N*_*ens*_ ≥ 200, our proposed method can adaptively predict the EEG time series and E/I ratio while overcoming the effects of nonstationary observation noise. These significant advantages of our proposed method are not seen in those previous conventional methods.

Despite the advantages of our proposed method, some limitations exist.

First, our proposed method can be applied only for single-channel EEG signal, that is, the proposed method cannot be applied to estimate the network-level dynamics of EEG signals. Even in the case of the parameter estimation for a single-column NM model from observed EEG, we must determine 11 state variables (six state variables and five model parameters) using the EnKF. Therefore, if we are to estimate the network-level dynamics of E/I balance changes from multiple-channel EEG data, an efficient parameter estimation algorithm with the EnKF scheme for high-dimensional data needs to be developed. In addition, when estimating the network-level dynamics of the E/I balance changes from multiple-channel observed EEGs, the effect of volume conduction should also be considered. To avoid such an effect, the application of current source density analysis in the preprocessing of the EEGs could be considered as a possible method for accomplishing this. However, to address these issues, further studies are required. Second, our proposed method remains the effect of the initial setting for state noise covariance *Q*. To avoid reducing estimation performance with nonstationary observed noise, the scale of observation noise covariance is recursively optimized using the variational Bayesian noise adaptation method. However, the scale of state noise covariance *Q* is fixed in our proposed method. To consider the effects of non-stationarity in both state and observed noise, the variational Bayesian algorithm we applied in this method should be modified. Third, in the neurophysiological validity evaluation of the current study, the EEG data that contained the awake period during the first NREM/REM cycle were excluded from the analysis because the effect of interrupted sleep on EEG oscillation and the analysis condition should be controlled in a comparison of prior evidence and our estimated results. As a result, the EEG analysis we conducted with our proposed method used a relatively small dataset. Therefore, in future work, we would like to conduct EEG analysis with our proposed method on a large dataset to discover the functional roles of the E/I balance changes.

## Methods

### Definition of the NM model

The NM model, proposed by Jansen and Rit [10], is a mathematical model for EEG that is formulated with the evolution of the average postsynaptic potential (PSP) in the three following interacting neural populations: pyramidal cells, excitatory interneurons, and inhibitory interneurons. This model can be described using the following six first-order differential equations [10]:

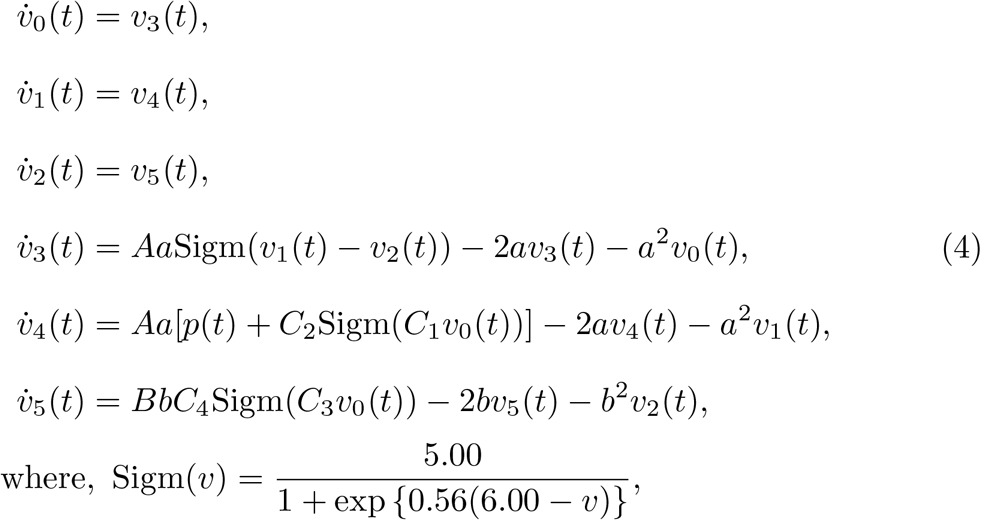

where variables *v*_0_, *v*_1_, and *v*_2_ represent the postsynaptic potential of pyramidal cells, excitatory interneurons, and inhibitory interneurons, respectively. The synthetic EEGs generated in the NM model are calculated by *y*(*t*) = *v*_1_(*t*) − *v*_2_(*t*). The fixed constants *C*_*n*_ (where, *n* = 1, …, 4) account for the number of synapses established between two neuron populations. In this study, *C*_*n*_ was fixed as *C*_1_ = 135, *C*_2_ = 108, and *C*_3_ = *C*_4_ = 33.75, drawing from previous studies [10]. The other five parameters in the NM model, *A, a, B, b* and *p*, are time-variant parameters and the target-to-state estimation in the vbcEnKF scheme based on observational EEG data in our proposed method. Parameters *A* and *B* represent the E and I synaptic gains, respectively. These two parameters control the amplitude of the postsynaptic potentials (PSP) generated by E and I interneurons. The parameters *a* and *b* indicate the inverse of the time constant for excitatory PSP (EPSP) and inhibitory PSP (IPSP). The rhythm of oscillation (dominant frequency band) of the EEG signals generated in the NM model is configured by *a* and *b*.

Parameter *p* represents the background noise input of the model. The behavior of the EEG signals generated in the NM model is bifurcated by the value of *p*. In this study, *p* was estimated within the range of 120–320, based on the study by Jansen and Rit (1995) [10] that showed that alpha-like limit cycle behavior was observed in the generated signal of the NM model when *p* was distributed in the above range. Moreover, an interval constraint for estimation of the inverse of the time constant *a* in EPSP was chosen as *a* = 5–200 to satisfy *τ*_*e*_[ms] = 5–200, where *τ*_*e*_ = *a*^*−*1^. Additionally, the constraint for the inverse of the time constant *b* in IPSP was chosen to satisfy *τ*_*i*_[ms] = 5–200, where *τ*_*i*_ = *b*^*−*1^. The boundary intervals for parameters *a* and *b* were selected so that the NM model could generate delta to beta rhythms, on the basis of the report by David and Friston (2003) [32].

### State and parameter estimation of the NM model

To track the time-varying changes in the NM model states and parameters, we applied the modified algorithm of an EnKF scheme. The EnKF is a well-known recursive data forecasting algorithm for data assimilation and is used in the research field of weather forecasting [8, 9]. Moreover, the EnKF works well for predicting time series of nonlinear dynamical systems. Since the observed EEG signal can be formulated using a nonlinear system such as the NM model, the EnKF scheme can be applied to directly estimate the system’s state and parameters in the NM model from the observed EEG data. However, the extent of the noise contained in the time series of observed neurophysiological signals such as EEG is usually unknown, and the noise component can be nonstationary. In addition, as we mentioned earlier, when estimating the NM model’s state and its five parameters *A, a, B, b* and *p* from observed EEGs, we gave the boundary constraint for each parameter so that the signal generated in the NM model showed EEG-like limit cycle behavior (see Supplementary Information for details). To estimate the state and parameters of the NM model from observed data while considering these issues, we combined a variational Bayesian noise adaptive algorithm in a linear/nonlinear Kalman Filter [12–16] and constrained state estimation in the Kalman Filter scheme [17–20] with a conventional EnKF [8, 9]. This method is hereinafter referred to as the variational Bayesian noise adaptive constrained EnKF (vbcEnKF). In this section, we will give a short description of the vbcEnKF algorithm for state and parameter estimation in the NM model. A more detailed mathematical description is given in the Supplementary information.

To apply the vbcEnKF approach to estimate the states and parameters in the NM model, the six first-order differential equations in the model were transformed to the state-space form, as shown below.

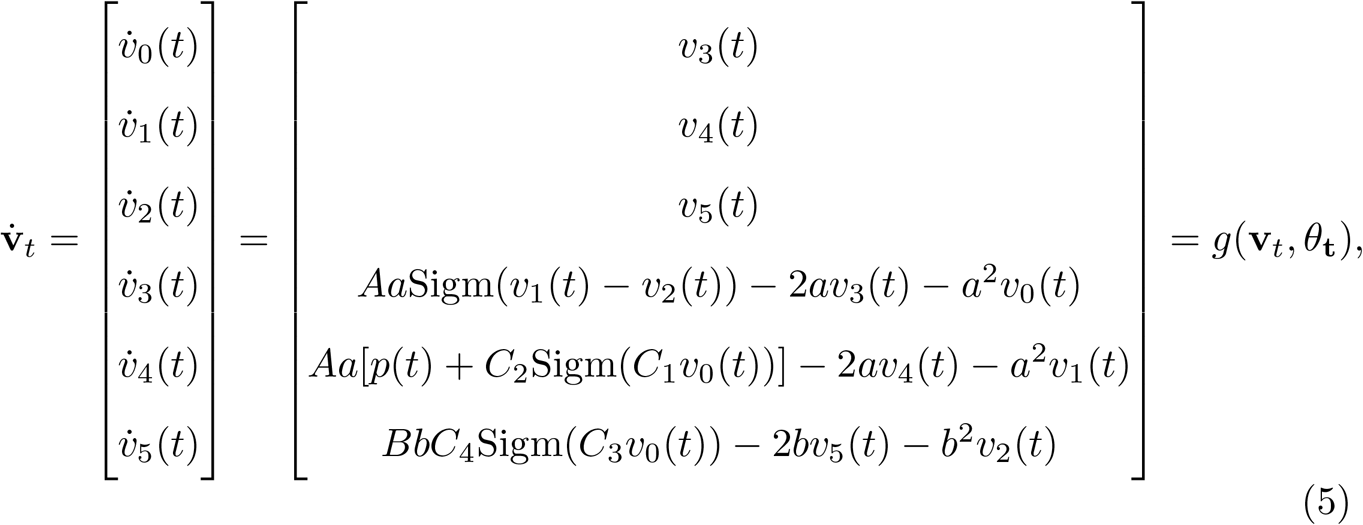

state model :

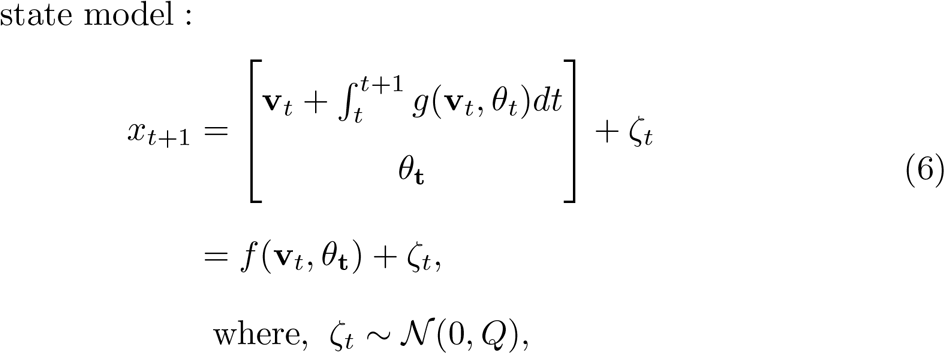

observation model :

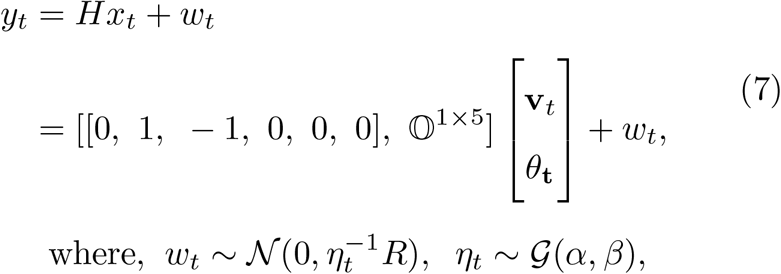

where **v**_*t*_ = {*v*_0_(*t*), *v*_1_(*t*), …., *v*_5_(*t*)} and *θ*_*t*_ = {*A*(*t*), *a*(*t*), *B*(*t*), *b*(*t*), *p*(*t*)} indicate the state and parameter vectors of the NM model, respectively. Note that 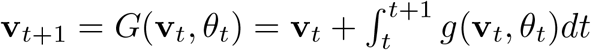 was implemented with the fourth-order Runge-Kutta method with step size Δ*t* (sampling interval). The variable *ζ*_*t*_ is state noise that follows the normal density 𝒩 (0, *Q*). Variable *w*_*t*_ is observation noise, which follows the normal distribution 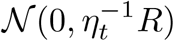.

Moreover, *η*_*t*_ is a noise scale parameter that follows the gamma probability 𝒢 (*α, β*). As mentioned above, applying the variational Bayesian scheme, the observation noise covariance 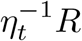 is adaptively optimized by estimating the noise scale parameter, *η*_*t*_, on a sample-by-sample basis [12–16]. See the Supplementary information for more details.

By applying the above-mentioned state-space form of the NM model to the vbcEnKF algorithm, model state and parameters were recursively estimated from observed EEG data on a sample-by-sample basis. The whole algorithm of this method includes the following five steps:

1. Initialize the parameters for the probabilities *x*_0_ ∼ 𝒩 (*x*_0_, *P*_0_), *ζ*_0_ ∼ *N* (0, *Q*), 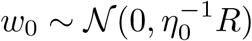, and *η*_0_ ∼ 𝒢 (*α*_0_, *β*_0_).
2. Predict the system’s state based on the current parameters of the probability density 𝒩 (*x*_*t*_, *P*_*t*_) in the state model (see the Prediction step in Algorithm 1)
3. Get new observed data 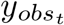 and update the parameters of probability densities of the state model and noise scaling factor *η*_*t*_ (see the Update step in Algorithm 1).
4. After updating the state *x*_*t*+1_, if the constraints for variable *x*_*t*+1_ are violated, the projection of the updated variable *x*_*t*+1_ in the Update step (see Algorithm 1) is corrected so that the variables are within a feasible range.
5. Repeat steps 2–4.

For step 1, the initial parameter value of the probability density for the state variable *x*_*t*_ was set at *x*_0_ = 𝕆 ^1*×*11^ (Note: *x*_*t*_ = {**v**_*t*_, *θ*_*t*_}, **v**_*t*_ ∈ ℝ^6^, *θ*_*t*_ ∈ ℝ^5^). The observation noise covariance *R* was set at *R* = 50 in both the numerical simulation and real EEG data analysis. The initial setting for 𝒢 (*a*_0_, *b*_0_) was set to *a*_0_ = 1 and *b*_0_ = 0.5. The state noise covariance Q was selected differently depending on the analysis. In the numerical simulation, the covariance *Q* was fixed as *Q* = diag ([Δ*t* ·10^*−*2^ ·𝕀^1*×*6^, 10^*−*3^ ·𝕀^1*×*5^]). In the EEG analysis, *Q* was fixed as *Q* = diag ([Δ*t* ·𝕀^1*×*6^, 10^*−*3^ ·𝕀^1*×*5^]). Δ*t* indicates that the sampling interval was set as Δ*t* = 0.01 in this study. The number of ensembles *N*_*ens*_ for the EnKF scheme was specified in each analysis (see the following section for more details). The summary of the whole algorithm of vbcEnKF is shown in Algorithm 1, and mathematical details are presented in the Supplementary information.

#### Algorithm 1

The variational Bayesian noise adaptive constrained Ensemble Kalman Filter (vbcEnKF) algorithm

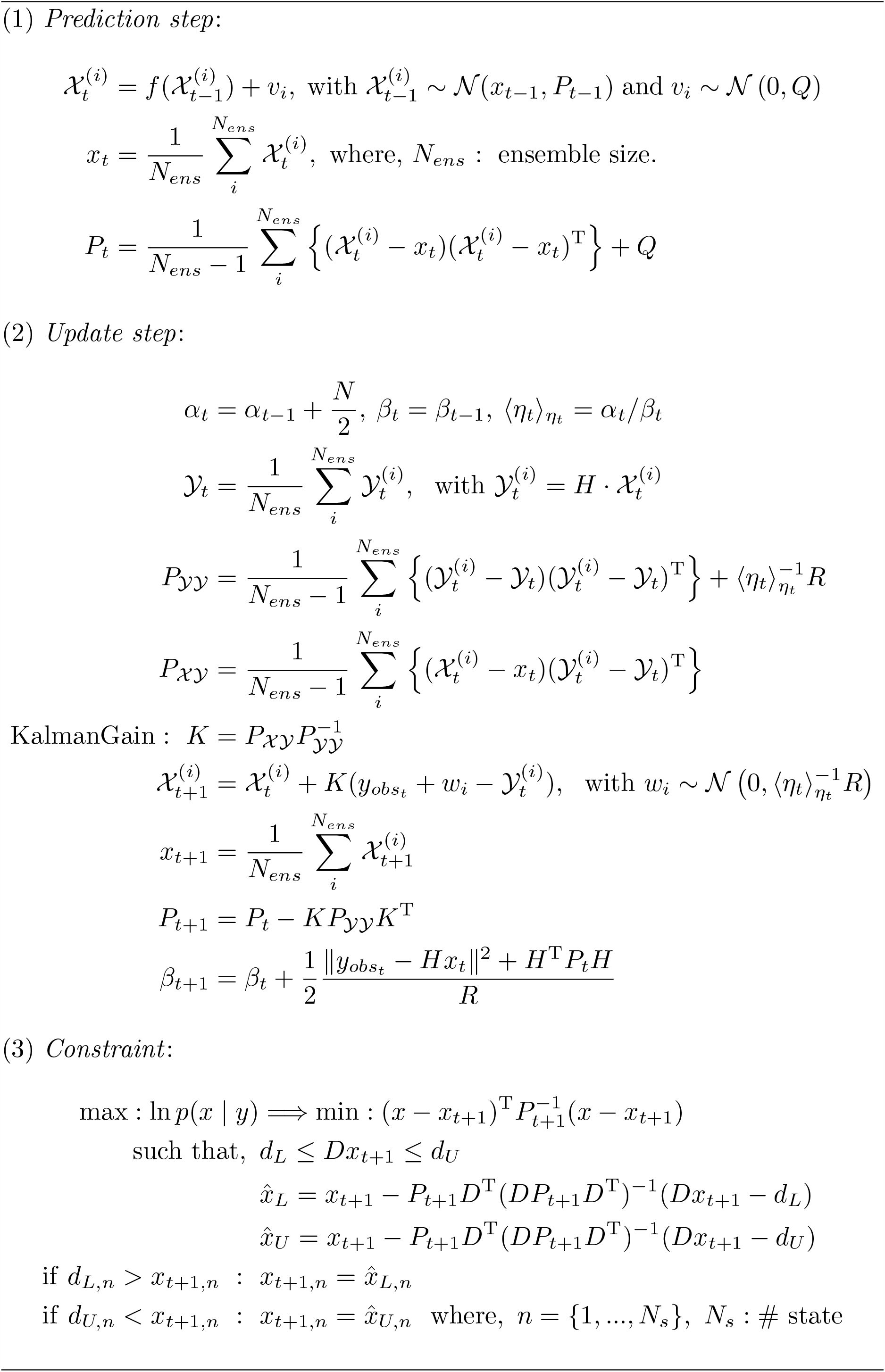

### Estimation of the mE/I ratio

As mentioned above, the amplitude of the EPSP and IPSP produced by the NM model can be influenced by the E and I synaptic gain parameters (*A, B*). These parameters play a key role in mediating the characteristics of the synthetic EEGs. Therefore, in our proposed method, *A* and *B* were estimated from observed EEG signals by the vbcEnKF scheme.

Subsequently, the E/I balance is evaluated as the temporal changes in the ratio of *A* to *B*. To track time-varying changes in this putative E/I ratio, we proposed the index of the E/I ratio, which we named the model-based E/I ratio (mE/I ratio), calculated using the following equation:

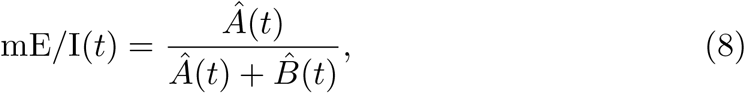

where *Â* (*t*) and 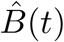 are E and I synaptic gain parameters estimated by our proposed method. The value of the mE/I ratio is sequentially calculated when updating the parameters in the NM model, on a sample-by-sample basis.

### Numerical simulation

To confirm the validity of our proposed method, we first applied our method to synthetic data generated by the NM model to clarify whether our method can sequentially estimate changes in the model parameters. Moreover, since the EnKF algorithm is based on the sMC method for Bayesian probability estimation, we considered the following two conditions under which to evaluate the effects of state and parameter estimation accuracy. First, we tested the effects of the ensemble size for the sMC method in the EnKF scheme. In this simulation, the number of ensembles selected ranged from 40 to 500, with 20 steps. Second, we tested the effects of the initial seed of the random generator in the sMC. Since this value would affect the time evolution of the state estimation in the NM model, our proposed method was applied to the synthetic EEG data 50 times with different initial random seed values for each ensemble size condition.

The synthetic EEG were generated by the NM model using a numerical integration with the fourth order Runge-Kutta method with the time step *dt* = 0.01 (i.e., a sampling frequency of 100 Hz). The total sample length of the generated data was 3, 000 (i.e., the total time length of each dataset was *t* = 30 s). The generated synthetic EEG data consisted of two segments separated by the event of the parameter changes. The exact value of parameters and the timing of the parameter changes are shown below.

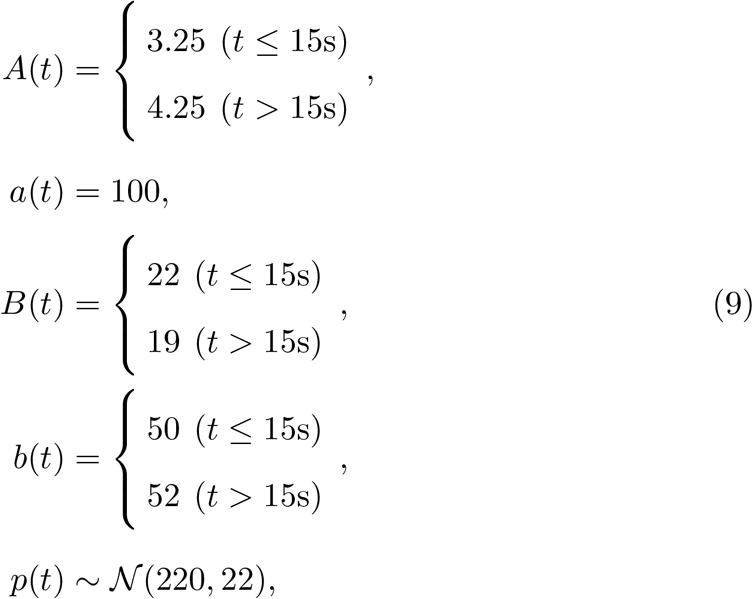

After generating synthetic EEG data with the above parameter settings, we applied our proposed method 50 times with a different initial random seed for each ensemble size condition. To test whether our method could correctly detect the parameter changes from observed data with noisy sampling, white noise was added to the synthetic EEG data using N(0, 1.3). The covariance of the generated white noise is set to *σ*^2^ = 1.3 to create a situation that considers the synthetic observed EEG signals to be strongly distorted by large observational noise. In this simulation setting, the synthetic observed EEGs are generated to satisfy the condition that the signal-to-noise ratio (SNR) is under 5.0 (i.e., log_10_(5) ≈ 0.699 dB). The exact SNR value of the synthetic observed EEGs was 3.838 (i.e., 0.598 dB).

Next, we evaluated the estimation error score of synthetic EEGs and the standard deviation of estimated noise covariance, which was obtained from 50 trials of the estimation for each ensemble size condition. The estimation error score was calculated using the mean absolute error function described by the following equation:

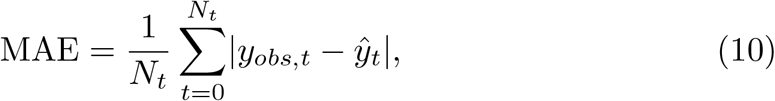

where *y*_*obs,t*_ and *ŷ*_*t*_ are the observed (exact) and predicted EEG at time sample *t*, and *N*_*t*_ is the total number of samples.

### EEG analysis

To confirm its neurophysiological validity, we applied our proposed method to an open EEG dataset [11] to assess whether it could detect changes in E/I balance between the NREM/REM sleep stages.

### Open EEG dataset

The open dataset used consisted of EEG data recorded with a bipolar montage and a single pair of electrodes located in Pz and Oz areas based on a 10–10 system (sampling frequency = 100 Hz). In this dataset, the EEG data were obtained from 197 participants in their own homes during day-night periods by Kemp et al. (2000) [11]. A more detailed description of the task settings is provided in the original paper. This dataset is available at the following PysioNet repository: https://physionet.org/content/sleep-edfx/1.0.0/#ref1. During this validation, datasets that contained awake periods during the first NREM period in sleep marker data were excluded to control the analysis conditions and avoid the effect of interrupted sleep on the EEG activity. As a result, we only used EEG data of 19 healthy participants who displayed a typical transition between sleep states during the first NREM and REM periods. Moreover, in the following validation, we used EEG data epochs that included the first 20-minute period from sleep onset time to the end of the first REM period for each participant.

### Preprocessing of EEG

To reduce the effects of artifacts (e.g., nose movement, eye movement, and eye blink signals), we first applied an artifact removal method with empirical mode decomposition [33, 34]. A detailed description of this method is provided in the original articles [33, 34]. After removing artifacts, EEG data were band-pass filtered between 0.6 and 20 Hz with a zero-phase finite impulse response filter (the number of taps = 6,000; transition with 0.1 Hz; window function = Hanning window).

### E/I ratio evaluation

After preprocessing, the EEG data were applied the vbcEnKF algorithm, and the mE/I ratio was sequentially calculated when updating the state and parameters of the NM model on a sample-by-sample basis. For the statistical analysis of the mE/I ratio, a temporally averaged mE/I ratio for the distinct five sleep stages (awake, stage 1, Stage 2, Stage 3/4, and REM) of each participant was calculated. The Kruskal-Wallis test was used to examine the differences in the mE/I ratio between the sleep stages. Moreover, to determine whether there were significant differences in the rank mean between specific pairs of sleep-stage-averaged mE/I ratios, Dunn’s multiple comparisons test (two-tailed test) was also applied after the Kruskal-Wallis test.

### Comparison with a prior method

To compare the estimations of sleep-dependent changes in the E/I ratio between our proposed method and prior methods, we applied the previous method of E/I estimation proposed by Gao et al. (2017) [23] to the same sleep EEG datasets [11] that we used in this study. The previous E/I estimation method [23] evaluated E/I balance changes using power spectral density (PSD) analysis of EEG signals. As mentioned in the Result and Discussion section, Gao et al. (2017) [23] found that the slope of the log-log plot of PSD from 30 to 50 Hz (i.e., 1/f power law exponent in the gamma frequency band) was correlated with the changes in the E/I ratio based on numerical simulation, which indicates that the E/I balance can be quantified according to this slope (hereinafter called the E/I slope method) [23]. More details on the E/I slope method can be found in the original article [23]. The sleep-dependent E/I balance changes for the EEG datasets used in this study were also evaluated by applying the E/I slope method, and these results were compared with those generated using our proposed method.

## Supplementary information

The Supplementary methods associated with this article can be found in the online version.

## Acknowledgments

This research was partially funded by the JSPS KAKENHI grant (20K19867, https://kaken.nii.ac.jp/en/grant/KAKENHI-PROJECT-20K19867/) from the Japan Society for the Promotion of Science, the NINS program of Promoting Research by Networking among Institutions (Grant Number 01412303), and the support for in-house research on data science approaches to physiological research at National Institute for Physiological Sciences, Japan. We would like to thank Assoc. Prof. Yoshihiro Noda and Dr. Masataka Wada for fruitful discussions and comments about TMS-EEG studies of the functional mechanism underlying E/I balance changes in human subjects.

## Data availability

The raw sleep EEG dataset obtained by Kemp et al. (2000) [11] is available at https://physionet.org/content/sleep-edfx/1.0.0/#ref1. All data used to reconstruct the results described in this paper are available at Github: https://github.com/myGit-YokoyamaHiroshi/EEG vbcEnKF mEI est.

## Code availability

The programming code used to reconstruct all the results in this paper was implemented using the language Python and is available at Github:

https://github.com/myGit-YokoyamaHiroshi/EEG vbcEnKF mEI est.

## Author contributions

**Hiroshi Yokoyama**: Conceptualization, Data curation, Formal analysis, Funding acquisition, Investigation, Methodology, Resources, Software, Validation, Visualization, Writing original draft, Writing – review & editing. **Keiichi Kitajo**: Project administration, Supervision, Resources, Validation, Writing – review & editing.

## Supplementary information

### S1. Variational inference for the Kalman Filter

As stated in the main manuscript, to consider the effects of unknown or nonstationary observation noise, we applied a variational Bayesian noise adaptive approach for the Kalman Filter (vbKF; Dong & Song, 2013; Sarkka & Nummenmaa, 2009; Stroud & Bengtsson, 2007; Wang & Guan, 2015; Wang et al., 2017). First, we will explain the mathematical details of the noise adaptive algorithm. For the sake of simplicity, we will here give a detailed description of the algorithm for the vbKF in a Gaussian linear case. A discrete-time linear state space model is defined using the following equations:

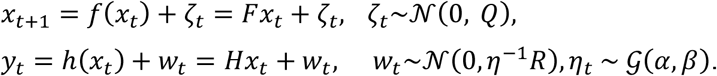

We here assumed that the probability of the observation noise follows the normal-gamma probability density. Furthermore, observation *y*_*t*_ and state variable *x*_*t*_ follow the multivariate Gaussian distribution 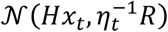 and 𝒩 (*m*_*t*_, *P*_*t*_), respectively. In such a case, if the state variable *x*_*t*_ and noise scaling parameter η are independent, the joint posterior probability density for these variables can be estimated with the variational Bayesian approximate, as below (Sarkka & Nummenmaa, 2009):

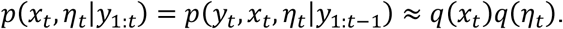

The variational Bayesian approximation can be solved by minimizing the Kullback–Leibler (KL) divergence between the approximated distribution *q*(*x*_*t*_)*q*(η_*t*_) and the exact posterior *p*(*y*_*t*_, *x*_*t*_, η_*t*_|*y*_1:*t*−1_), as below:

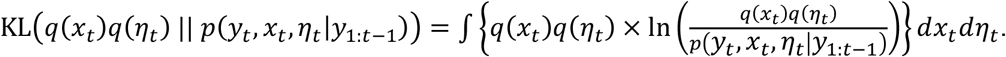

Minimizing the right term of the above equation with respect to the probability *q*(*x*_*t*_), the solution of updating rule of the probability density for *x*_*t*_ is given as the following equation:

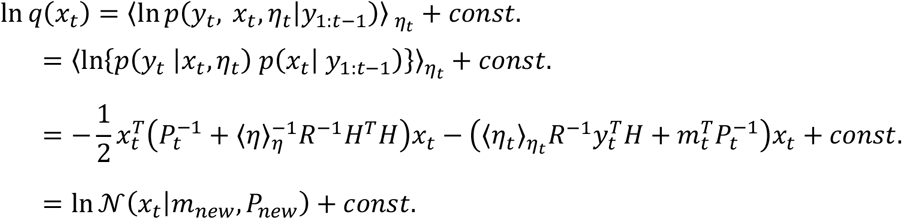

This means that the parameters *m*_*new*_ and *P*_*new*_ can be calculated as follows:

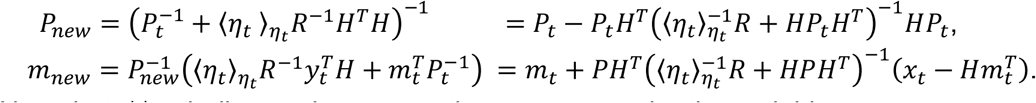

Note that 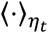 indicates the expectation operator under the variable η_*t*_.

In the same manner, the solution of *q*(η_*t*_) is also given, as follows:

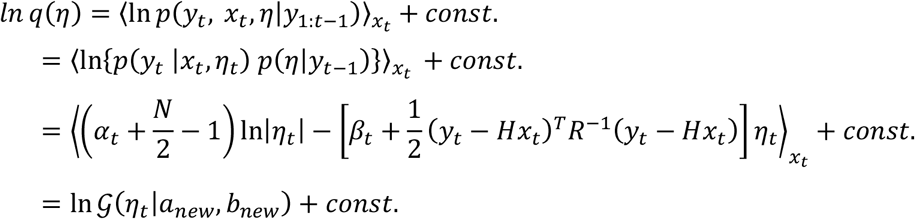

Note that 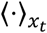 indicates the expectation operator under the variable *x*_*t*_. The parameters *a*_*new*_ and *b*_*new*_ can be calculated as follows:

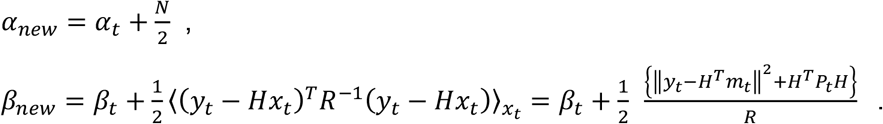

The expectation of parameter η_*t*_ (i.e., 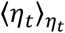) can be described as follows:

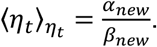

### S2. Inequality constraint for Kalman Filter

In this section, we will explain the details of the state estimation in KF with an inequality constraint. In the context of the study on the control theory in robotics, various approaches of imposing the constraint in state estimation for the KF scheme have been proposed (Simon & Chia, 2002; Simon, 2010). Among these, we chose to apply the estimation projection approach with inequality constraints (Luzar et al., 2012; Simon & Chia, 2002; Simon, 2010; Wang et al., 2009) in our proposed method. In this approach, if the state variables *x*_*t*_ follow the probability density *x*_*t*_ ∼ *p*(*x*_*t*_|*x*_*t*−*t*_) = 𝒩 (*m*_*t*_, *P*_*t*_) with the constraint *Dx*_*t*_ ≤ *d*_*U*_, the KF algorithm estimates the value of *m*_*t*_ and *P*_*t*_ so as to solve the following problem:

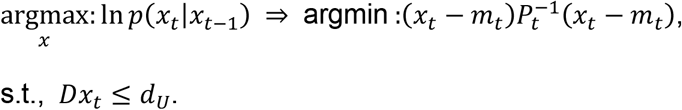

This problem can be solved using a Lagrange multiplier method, as follows:

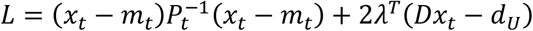

The solution is given as follows:

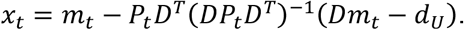

In the same manner, we can solve the solution in the case of the constraint *Dx*_*t*_ ≥ *d*_*L*_, as follows:

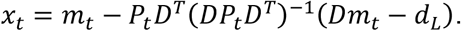

In the current study, to estimate the state and parameter using the EnKF scheme while considering parameter constraints so as to generate the periodic signal in the NM model, we applied the following interval constraint:

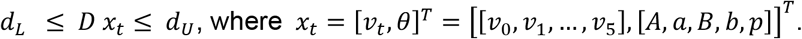

Note that parameters *D, d*_*L*_, and *d*_*U*_ followed with *D* = [𝕆^5×6^ 𝕀^5×5^], *d*_*L*_ = [0.01, 5.00, 0.01, 5.00, 120]^*T*^, and *d*_*U*_ = [100.00, 200.00, 100.00, 200.00, 320.00]^*T*^; that is, such interval constraints were set to restrict the range of values in the model parameters *A, a, B, b*, and *p*. In particular, interval constraints for the parameter *p* in the NM model were selected so that the estimated value of *p* was satisfied with the interval 120 ≤ *p* ≤ 320. This was chosen based on the study by Jansen and Rit (1995), which showed that the alpha-like periodic simulated signal was observed in the NM model when the external input *p* was distributed in the range of 120 ≤ *p* ≤ 320. An interval constraint for the inverse of time constant *a* in excitatory post-synaptic potential (PSP) was chosen as 5 ≤ *a* ≤ 200 to satisfy 5 ≤ τ_*e*_[ms] ≤ 200, where τ_*e*_ = *a*^−1^. Also, the constraint for the inverse of time constant *b* in inhibitory PSP was chosen to satisfy 5 ≤ τ_*i*_[*ms*] ≤ 200, where τ_*i*_ = *b*^−1^. The parameters *a* and *b* were selected so that the NM model can generate delta to beta rhythms based on the report by David and Friston (2002).

### S3. Variational Bayesian noise adaptive constrained Ensemble Kalman Filter (vbcEnKF)

S1 and S2 described the whole algorithm of the proposed vbcEnKF. To estimate state and model parameters in the NM model, we combined the variational Bayesian noise adaptive algorithm and inequality constraint approach with the EnKF. The whole vbcEnKF algorithm is summarized in the following table.

**Supplementary Table 1:**
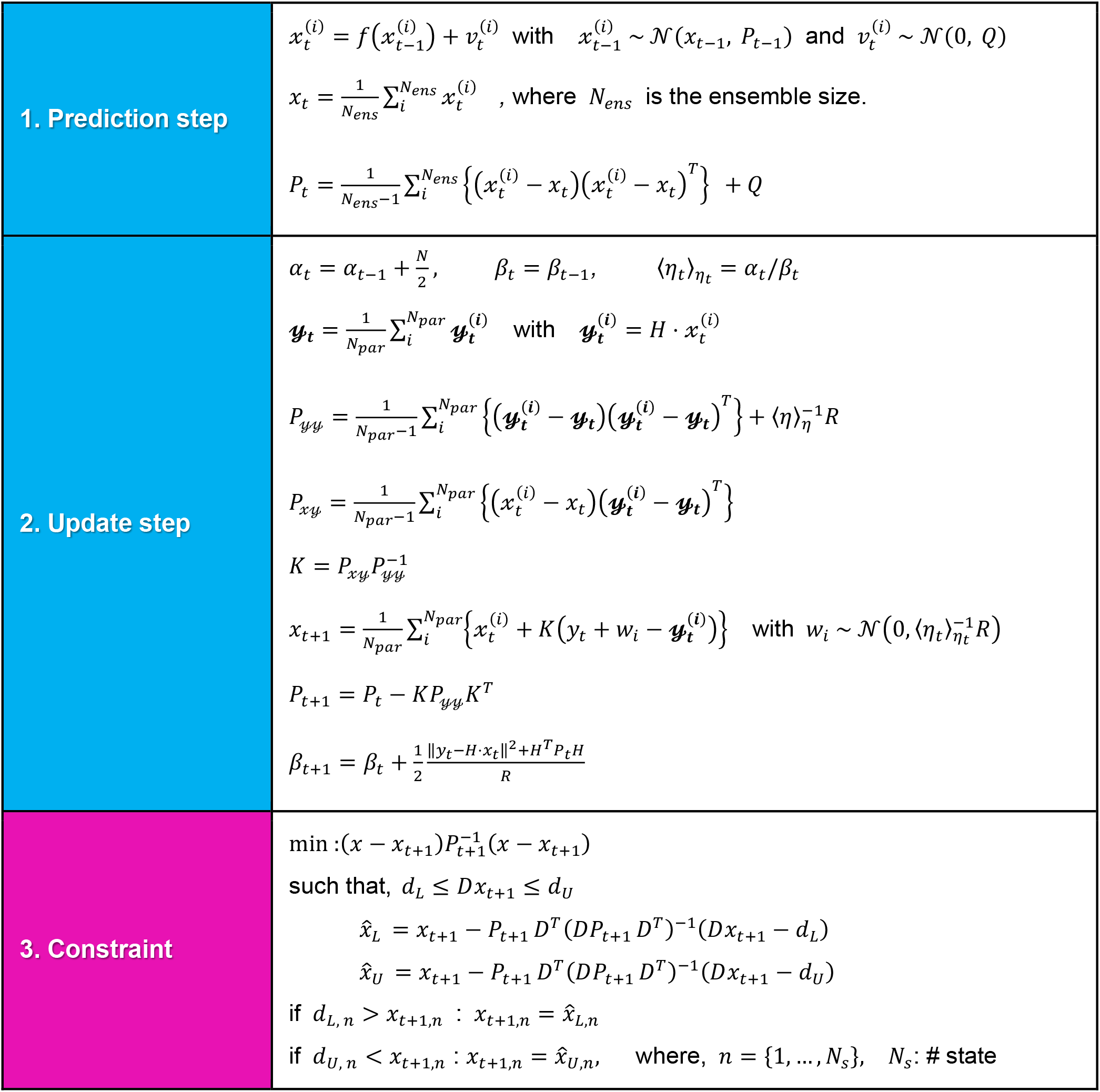
Algorithm of the vbcEnKF. The vbcEnKF consists of three steps. The first step is predicting the state variables and parameters in the NM model based on the current probability in the estimated state model. The second step is model updating the probability in the state and the observed model using observed data. The third step is applied only if the constraint for the updated state variables is violated.

